# ntSynt-viz: Visualizing synteny patterns across multiple genomes

**DOI:** 10.1101/2025.01.15.633221

**Authors:** Lauren Coombe, René L. Warren, Inanc Birol

## Abstract

With the explosion of chromosome-scale genome assemblies being generated in recent years, there is vast potential for comparative genomics analyses through detecting multi-genome synteny. While existing tools can detect synteny blocks between multiple genomes, their text-based outputs make it challenging to intuitively explore large-scale synteny patterns. Interpretable, information-rich and easy-to-use synteny visualization tools are imperative to enable important biological insights from the synteny block data output by the aforementioned utilities. Here, we present ntSynt-viz, a command-line tool for automated sorting, normalization and plotting of multi-genome synteny blocks. We show how ntSynt-viz provides clearer and more easily interpretable chromosome-painting ribbon plots compared to the state-of-the-art tool NGenomeSyn when evaluating synteny between 14 human genomes and 9 hoverfly genomes. We expect that ntSynt-viz will provide crucial insights into large-scale synteny patterns between divergent genomes, thereby advancing research into key evolutionary questions. ntSynt-viz is freely available on GitHub (https://github.com/bcgsc/ntsynt-viz).

## Introduction

Studying genome synteny, the conservation of genome structure, across multiple genomes allows for important insights into the evolution of both species and populations (Liu, Hunt and Tsai 2018). Understanding how large-scale genome structures have evolved over time through rearrangements and inversions helps researchers to reconstruct the evolutionary history of the genomes, and thus helps better understand the functional impacts of those changes (Ghiurcuta and Moret 2014). With an increasing number of chromosome-level genomes being assembled by the community, particularly through large consortia like the Human Pangenome Project and the Earth BioGenome Project, there is immense potential to leverage this vast and diverse data to uncover important evolutionary insights (Lewin *et al*. 2022; Wang *et al*. 2022).

A number of bioinformatics tools detect synteny between whole genome assemblies, as opposed to only using a restricted number of homologous genes, for example. Most of these whole-genome synteny tools, such as SyRI (Goel *et al*. 2019), halSynteny (Krasheninnikova *et al*. 2020), and Satsuma (Grabherr *et al*. 2010), are limited to pairwise comparisons between genomes. Other tools, such as ntSynt, the computationally lightweight synteny detection utility we developed (Coombe *et al*. 2024), can identify synteny between multiple genomes simultaneously, which allows for more flexible and rich comparisons between divergent genomes. However, the output files generated by these synteny tools are generally structured text files which, though information-rich, may be difficult for users to interpret in isolation. Therefore, it is essential to develop a streamlined, user-friendly and aesthetically pleasing visualization utility to allow researchers to gain insights on genome conservation by clearly depicting any syntenic relationships between sequences of interest.

Currently, there are a number of command-line tools and R packages for visualizing synteny blocks, each of which vary in their flexibility and ease of use. NGenomeSyn (He *et al*. 2023) and plotsr (Goel and Schneeberger 2022) are both command-line tools that generate ribbon plots to visualize computed synteny blocks, and both can plot multiple (i.e., more than 2) genomes. Both allow for synteny block coordinates to be specified in conventional formats, although plotsr is tightly coupled with the pairwise synteny detection tool SyRI (Goel *et al*. 2019). NGenomeSyn allows for more flexible layouts and visualization of inter-chromosomal rearrangements, while the inter-chromosome synteny visualization option for plotsr is marked as experimental. While R packages such as gggenomes (Hackl, Ankenbrand and Adrichem 2023) and syntenyPlotteR (Quigley *et al*. 2023) can also generate visualizations from detected synteny blocks, they require fairly extensive file formatting and pre-processing to adhere to their input requirements, as well as knowledge of the R programming language, which may be less accessible to some researchers. None of these existing tools or packages offer in-code pre-processing of the synteny blocks to allow for a clear and intuitive visualization. Thus, there is an unrealized need for an easy-to-use synteny visualization tool with built-in pre-processing steps to produce informative and easily understandable representations of syntenic relationships between multiple genomes.

Here, we present ntSynt-viz, a command-line visualization tool that generates publication-grade chromosome painting ribbon plots from detected synteny blocks. Powered by a single command, ntSynt-viz incorporates multiple important new features, such as leveraging synteny block mappings to 1) order the input chromosomes based on structural similarity, 2) normalize the strands of the input chromosomes compared to a target genome, and 3) utilize synteny-based distance estimations for top-to-bottom ordering of the genomes. In addition, using gggenomes (Hackl, Ankenbrand and Adrichem 2023), we integrate chromosome painting-inspired (Ried *et al*. 1998) colouring with ribbon plots to further enhance the interpretability of the output images. While ntSynt-viz works directly with synteny blocks computed by ntSynt (Coombe *et al*. 2024), it can also handle synteny blocks from other tools. We expect that the aesthetics provided by ntSynt-viz will allow for rich biological insights into the evolutionary history of diverse species and populations.

## Methods

ntSynt-viz enables user-friendly and informative depictions of multi-genome synteny blocks computed by ntSynt. To visualize synteny blocks computed by a different synteny detection tool, the synteny blocks simply need to be converted to the straightforward BED-like ntSynt format; we provide conversion scripts for multiple tools such as SyRI (Goel *et al*. 2019) and halSynteny (Krasheninnikova *et al*. 2020). The tool is run with a single python command, and is driven by a snakemake pipeline for ease of use.

### Genome sorting

To leverage the detected genomic synteny to sort the input genomes in the final ribbon plot, ntSynt-viz first estimates pairwise distances between each genome using the computed synteny blocks. To compute the distances, the consistencies between adjacent synteny blocks are assessed for each pair of genomes. To compute the estimated synteny distance between genomes *x* and *y* using *n* ordered multi-genome synteny blocks, we use Equation (1).

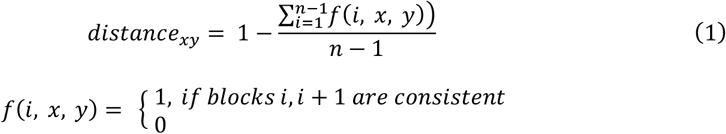

For synteny blocks *i* and *i* + 1 to be consistent in genomes *x* and *y*, they need to have the same strand transition (i.e. + **→** +; + **→** -, etc.), the start positions of blocks *i* and *i* + 1 need to be consistently increasing or decreasing, the chromosomes for each genome need to be the same between blocks *i* and *i* + 1, and the blocks cannot be separated by an indel (parameter *--indel*). After computing the pairwise distances, a neighbour-joining cladogram is constructed using Quicktree (Howe, Bateman and Durbin 2002), and the leaf ordering of this cladogram determines the genome ordering.

Optionally, a user can provide any newick-formatted cladogram using the *--tree* option, and the genomes will be sorted based on the leaves of this cladogram instead.

Should a particular genome be desired at the top of the final ribbon plot, users can specify it using *--target-genome*. In this case, the constructed or supplied cladogram will be rotated at nodes from the leaf to the root node, retaining the topology, but flipping the desired genome to the top. Starting from the first internal node parent of the target genome leaf, for each subtree of the given node moving towards the root, if the target genome is not at the top of the subtree, we perform a flip operation to move it to the top (Supplementary Figure S1). Otherwise, the subtree structure is retained. By iteratively performing this operation with increasingly deep subtrees that include the target genome, the desired genome is placed at the top of the cladogram, which will then ensure that it is also on the top of the final ribbon plot.

### Chromosome sorting

Similar to the top-down sorting of the genome assemblies, the chromosomes of the input genomes will be sorted from left to right based on the detected synteny blocks (Supplementary Figure S2). The chromosome sorting is performed on consecutive pairs of genomes, with the ordering determined as previously described. For each consecutive pair of genomes, *i* and *i* + 1 (0 <= *i* < *num_assemblies* − 1), each chromosome of genome *i* is split into tiles of length *-- tile* (default 1 Mbp). For each tile, the overlapping synteny blocks in genome *i* + 1 are tallied, building up a data structure which tracks the total lengths of each overlapping chromosome in genome *i* + 1. Then, the ‘best hit’ for that tile is chosen as the chromosome in genome *i* + 1 which has the largest total overlap length. This is repeated for all tiles across all chromosomes in genome *i*, yielding an ordered list of ‘best hit’ tiles. From that list, consecutive hits to the same chromosome in genome *i* + 1 are grouped together, and their lengths summed. Then, the list is filtered to only retain the representative chromosome entry which has the highest length. This list then dictates the ordering of chromosomes in genome *i* + 1.

### Strand normalization

Because independent genome assemblies can assemble structurally similar chromosomes on different strands, this feature leverages the detected synteny to normalize chromosome strands based on a target genome, taken as the first in the previously determined genome order, or the genome specified by *--target-genome*, if applicable.

For this normalization, for each chromosome of genome *x* (0 < *x* < *num_assemblies*), where genome 0 is the target, for each synteny block *j*, we determine if the coordinates of *j* have the same or different strands in genomes 0 and *x* . Once we have tallied the total synteny block lengths for the same (+) and different (-) orientations for all chromosomes, each chromosome of genome *x* is assigned to the orientation with the largest tallied length. These orientations are printed to a TSV file, and the synteny blocks reverse-complemented prior to plotting when the orientation is determined to be different (-). This feature is optional, and is activated by adding the *--normalize* option to the ntSynt-viz command.

### Ribbon plots

Using the files generated by the genome sorting, chromosome sorting, and strand normalization steps, gggenomes (Hackl, Ankenbrand and Adrichem 2023) is used to generate the chromosome painting ribbon plots, where the genomes are ordered top-to-bottom, and the chromosomes are plotted left-to-right. The ribbons are coloured based on the chromosome identity of the top-most (target) genome on the plot. Chromosome painting of each sequence is achieved by colouring each chromosome segment covered by a synteny block with the same colour scheme used for the ribbons. As the underlying chromosome is gray, regions that are not covered by a synteny block will be shown in gray. Optionally, a BED-like file with coordinates of gaps or centromeres can be supplied, and these will be plotted as black segments on the chromosomes (*--centromeres*). In addition, if a newick-formatted tree file is supplied to the program (using the previously described *--tree* option), this will be plotted next to the ribbon plot using ggtree (Yu *et al*. 2017). In addition to these features, the plots can also be customized in their dimensions (*--width, --height)*, the length of the scale bar (*--scale)*, the output format (*-- format png/pdf)*, minimum chromosome length to display (*--seq_length*), user-specified conversions of file names to names on the plot (*--name_conversion*), a subset of chromosomes to plot (*--keep)*, as well as the minimum synteny block length (*--length*). Finally, if different haplotypes of a given genome are being compared within the ribbon plot, the user can specify these relationships using *--haplotypes*, which will ensure that haplotype assemblies are adjacent and nudged together in the ribbon plot. Full explanations of all parameters for ntSynt-viz, including expected formats for the input files, can be found in Supplementary Table S1.

### Data

To demonstrate the functionalities of ntSynt-viz, we first ran ntSynt on two sets of genomes: 14 human genome assemblies and 9 hoverfly genomes (accessions listed in Supplementary Tables S2 and S3). Each human genome compared is chromosome-scale, and the genome assemblies were filtered to retain only the autosomes. The 14 human sequence assemblies come from 10 individuals, due to 5 individuals having diploid assemblies.

### Testing ntSynt-viz and NGenomeSyn

For each set of genomes, ntSynt v1.0.2 was run with default parameters to generate the synteny blocks, with the exception of specifying *-d* 0.5 and *-d* 20 for the human and hoverfly comparisons, respectively. To visualize the synteny blocks, ntSynt-viz v1.0.0 was run using these synteny blocks, with the commands launched shown in Supplementary Table S4. NGenomeSyn v1.41 was run using manually customized configuration files, which can be found on at https://doi.org/10.5281/zenodo.14247722.

## Results and Discussion

To demonstrate the utility of ntSynt-viz, we generated chromosome painting ribbon plots from two multi-genome synteny comparisons computed using ntSynt: comparing 14 human genome assemblies from 10 individuals, and 9 hoverfly genomes.

We first demonstrate the use of ntSynt-viz in comparing the 14 human genome assemblies (Figure 1). In the ribbon plots, following a given ribbon from the top to the bottom of the plot represents a multi-genome synteny block, with the colouring of the ribbon being dictated by the chromosome in the “target” (top) genome. The detected synteny is further highlighted with the use of the chromosome painting, where the regions that intersect with a given synteny block are coloured based on the incident synteny ribbons. Optionally, any user-specified regions can be plotted on the chromosome rectangles in black; we plotted the annotated T2T genome centromeres in Figure 1.

**Figure 1.**
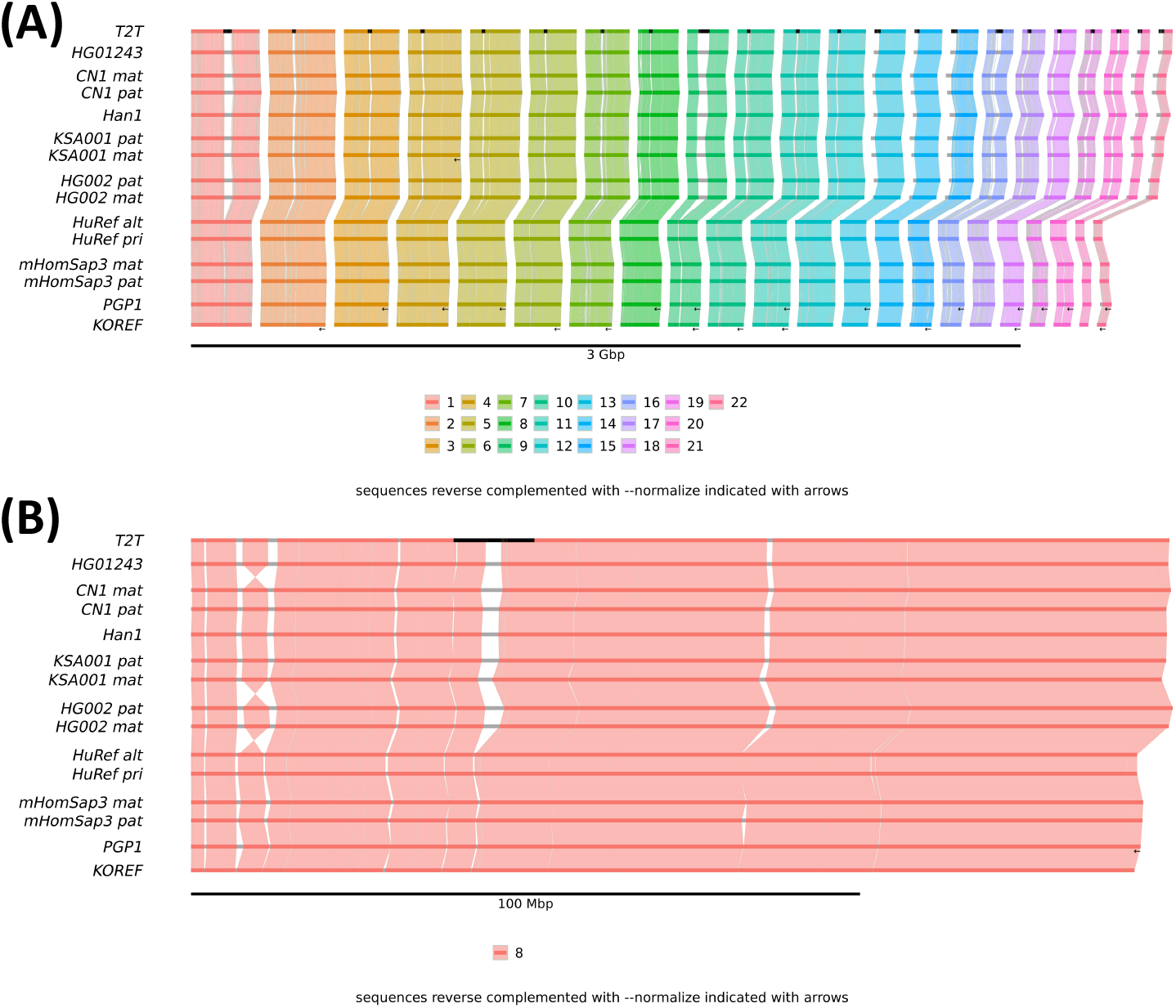
ntSynt-viz plots comparing 14 genome assemblies from 10 human individuals. (A) Plotting synteny blocks between all autosomes. The ribbons and chromosomes are coloured based on the chromosome of the corresponding synteny block in the T2T (target) genome at the top, and, when applicable, the black arrows below the rectangles indicate chromosomes that were reverse complemented by the strand normalization feature. The black bars on the T2T chromosomes represent the annotated centromeres. The scale for the chromosome lengths is indicated by the bar and label under the plot. (B) Zooming into the synteny blocks on chromosome 8.

As evident from the largely parallel and non-crossing ribbons, the human genome assemblies are largely syntenic, as expected (Figure 1A). The similarities between haplotypes of the diploid genomes are also highlighted by haplotype genome assemblies being nudged together in the visualization, distinguishing them from genomes from other individuals. The synteny block strand normalization also contributes to the clear synteny patterns, as 1, 12 and 9 chromosomes from KSA001 mat, PGP1 and KOREF, respectively, were reverse complemented to keep the chromosome strands consistent with the T2T reference genome. Full chromosomes being assembled on opposite strands is attributable to technical details of the genome assembly process and does not have any biological relevance, so performing this reverse-complementation enhances the clarity of the visualization, as is evident by comparing the ntSynt-viz plot to the plot generated by NGenomeSyn (Supplementary Fig S3).

The plot also reveals inversions near the beginning of chromosome 8, which are more clearly visualized when zooming into chromosome 8 for all genomes using the *--keep* option in ntSynt-viz (Figure 1B). In this zoomed-in view, there are clear twisting ribbons between HG01243 and CN1 mat, KSA001 mat and HG002 pat, and HG002 mat and HuRef alt. These top-to-bottom twisting patterns indicate that the inverted region is shared in the T2T, HG01243 and HG002 genomes. Interestingly, this region is a known polymorphic inversion in the human population (Feuk 2010; Logsdon *et al*. 2021).

To show the value of using ntSynt-viz to uncover synteny patterns between genome assemblies of non-model species, we detected 9-way synteny blocks between genome assemblies from the hoverfly genus *Cheilosia* using ntSynt, and visualized them using ntSynt-viz. These genomes were obtained from the Earth BioGenome Project database but were originally sequenced by the Darwin Tree of Life project (https://www.darwintreeoflife.org/). They vary in both chromosome number (5–7) and genome size (354–545 Mbp) (Supplementary Table S3) (Falk *et al*. 2021, 2023, 2024a, 2024b, 2024c; 2024d, 2024e, Crowley *et al*. 2023; Mitchell *et al*. 2024). As shown in Figure 2, there are large segments of the genomes that show considerable synteny, but also a number of intra-chromosomal inversions and differences in chromosome structure among the genomes. In particular, all compared genomes retain similar genomic content for the left-most plotted chromosome (red, OX411880.1 in *C. impressa*), but with a number of intra-chromosomal rearrangements and inversions. In addition, the bulk of the regions without detected synteny (seen in gray) also fall near the center of these chromosomes, suggesting the sequences are unique in at least one of the genomes, and perhaps partly due to genomic expansions in this region, as the genomes do vary up to ∼200 Mbp in size. These insights were enabled by the automated chromosome sorting, as the chromosome names provided in the input genomes were simply accession numbers, which themselves did not have any inherent order. Therefore, by automatically sorting the chromosomes using the synteny-driven approach implemented in ntSynt-viz, these synteny relationships became more apparent.

**Figure 2.**
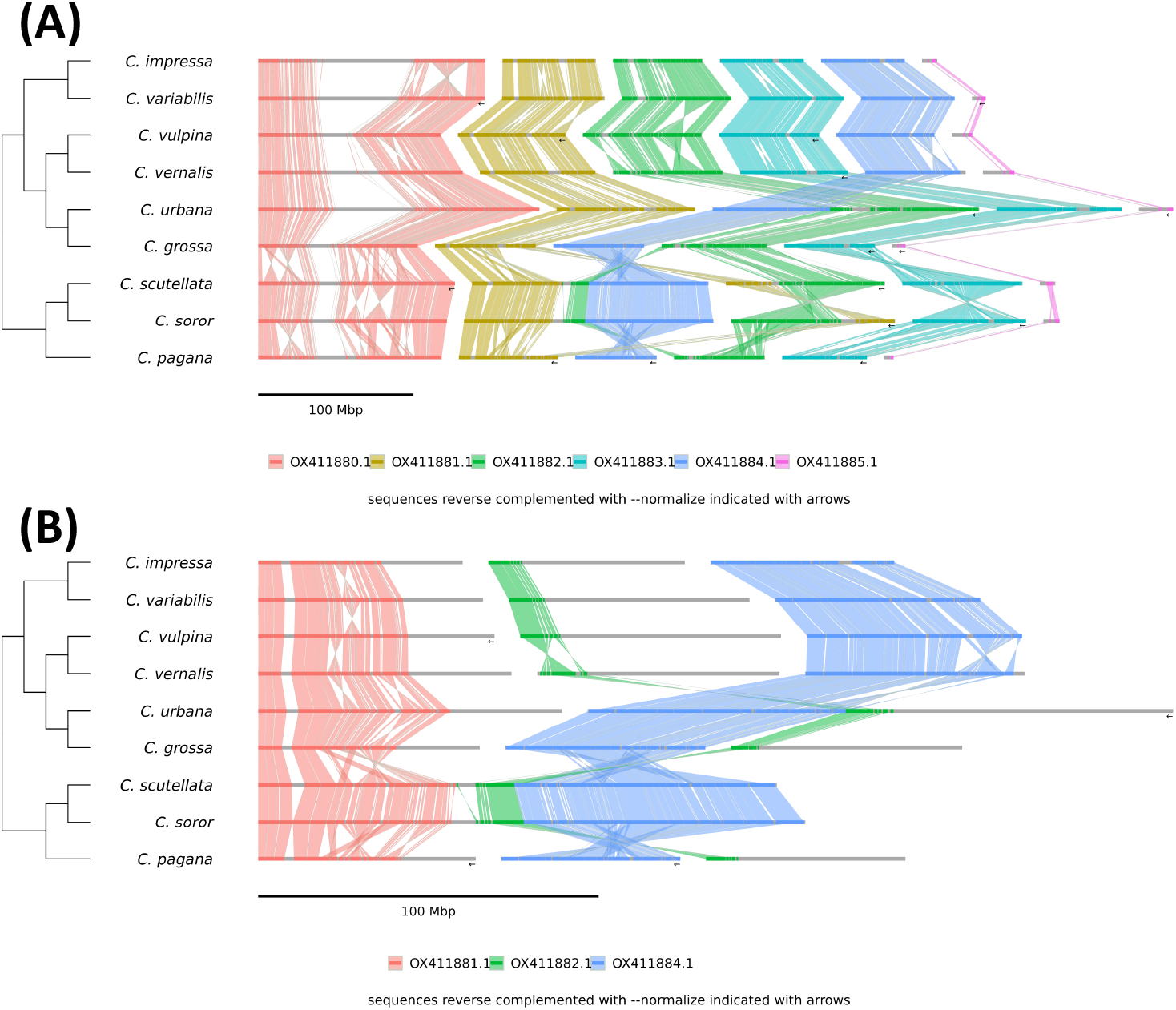
ntSynt-viz plots visualizing ntSynt multi-genome synteny blocks from 9 *Cheilosia* (hoverfly) genome sequence assemblies. (A) Plotting synteny blocks from all chromosomes. (B) Zooming into the synteny blocks that comprise the *C. soror* chromosome fusion.

Similar to the human plot, normalizing the strands of the chromosomes to the target (top) genome in the plot also aids in the clarity of the depicted rearrangements. For example, compared to the same ntSynt-viz plot without strand normalization (Supplementary Figure S4), for the chromosomes coloured yellow (OX411881.1 in *C. impressa*), the normalization of this chromosome in *C. vulpina* allows the synteny between both *C. variabilis* and *C. vernalis* to be more apparent. In the normalized version, observations including a couple of large inversions between *C. variabilis* and *C. vulpina*, and a high degree of collinearity between *C. variabilis, C. vulpina* and *C. vernalis* are clearer, while these observations are non-obvious in the non-normalized version. In addition to aiding in plotting a clearer picture of synteny, this feature can also be generally useful for normalizing strands between non-model genome assembly files; a list of the reverse-complemented chromosomes is available in an ntSynt-viz output file.

Interestingly, both *C. scutellata* and *C. soror* genomes have the most structural rearrangements as compared to the other genomes, including a particular fusion that is shared between these two genomes. This shared fusion is easily visualized due to the automated synteny-informed top-to-bottom sorting of the genomes, which highlights the similarities between closely related genomes, while still conveying differences between those that are more distantly related. In Figure 2B, we zoom into the synteny blocks that fall on this shared fusion, and see that it is a rearrangement of segments from 2 chromosomes in *C. urbana* and 3 different chromosomes in the other genomes. Interestingly, the majority of the blue ribbons (OX411884.1 in *C. impressa*), represent a full, individual chromosome in all genomes except *C. scutellata, C. soror* and *C. urbana*. Further research into these fusions, as well as other observable large structural rearrangements, would help to further uncover the evolutionary history of the *Cheilosia* genus.

In comparison, when visualizing the same *Cheilosia* ntSynt synteny blocks using NGenomeSyn, the synteny patterns and chromosome fusions are harder to interpret (Figure 3). While NGenomeSyn uses a similar ribbon plot approach, it does not have comparable pre-processing features, leading to plots that are less understandable, particularly for more complex syntenic relationships. In particular, the lack of automatic chromosome sorting means that the chromosomes, such as OX411880.1 in *C. impressa*, that have largely conserved content across the genomes are not easily seen. While in the ntSynt-viz plots, crossing ribbons more readily point to structural differences, for the NGenomeSyn plots, crossing ribbons may simply be due to the arbitrary ordering of the chromosomes. In addition, due to the random order of genomes top-to-bottom, the interesting shared chromosome fusion in *C. scutellata* and *C. soror* that is readily apparent in the ntSynt-viz plot is not easily discernable in the NGenomeSyn visualization. Importantly, manual scripts and configuration file adjustments were required to generate the figure, as opposed to the single command that drives ntSynt-viz. While NGenomeSyn ran faster than ntSynt-viz for visualizing the synteny blocks (0.1 min vs. 1.0 min for NGenomeSyn and ntSynt-viz, respectively), both tools finished in less than 2 minutes across all runs (Supplementary Table S5).

**Figure 3.**
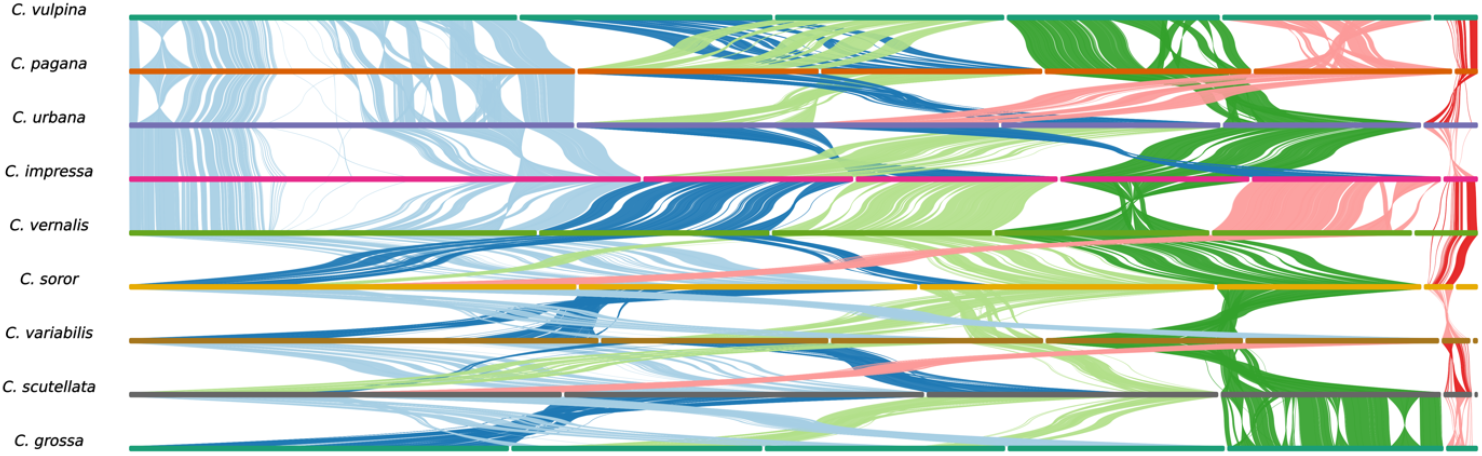
NGenomeSyn plots visualizing ntSynt multi-genome synteny blocks detected between 9 *Cheilosia* (hoverfly) genome sequence assemblies.

As demonstrated, the new sorting and normalization features introduced by ntSynt-viz allow for automated and clear visualization of syntenic relationships between multiple genomes, which can inform any number of downstream analyses. While other tools, such as NGenomeSyn, plotsr and SyntenyPlotteR, do provide the ability to visualize syntenic relationships as similar ribbon plots, they lack any automated synteny block pre-processing, meaning that these operations would have to be done manually by the user, which can be cumbersome and technically difficult.

In addition to the automated sorting and normalization features, ntSynt-viz is flexible to various customizations of the generated ribbon plots, with all options easily tweaked using command-line parameters. For example, the genome at the top of the plot, which dictates the chromosome and ribbon colouring scheme, can be specified using `--target-genome`. Furthermore, if using the tool with more fragmented genome assemblies, shorter sequences can be filtered out using --seq_length, and only synteny blocks above a given length can be plotted using --length. This filtering can help to produce clearer synteny visualization plots, as visualizing many small sequences can overcomplicated the general synteny patterns. The dimensions and output format of the plot itself can also be easily changed using command-line parameters.

The streamlined, user-friendly and automated visualizations of multi-genome synteny blocks enabled by ntSynt-viz will help researchers to decipher the interesting biological and evolutionary information inherent in the synteny blocks. These insights can enable deeper understanding of the evolutionary history of species, and help propel the progress of comparative genomics research to keep pace with the rapid increase in available genome assemblies.

## Code Availability

ntSynt-viz is freely available on GitHub (https://github.com/bcgsc/ntsynt-viz).

## References

Coombe L, Kazemi P, Wong J et al. Multi-genome synteny detection using minimizer graph mappings. bioRxiv 2024:2024.02.07.579356.

Crowley LM, Sivell O, Ashworth M et al. The genome sequence of the Figwort Cheilosia, Cheilosia variabilis (Panzer, 1798). Wellcome Open Res 2023;8:377.

Falk S, Crowley LM, Poole O et al. The genome sequence of the Truffle Blacklet, Cheilosia soror (Zetterstedt, 1843). Wellcome Open Res 2023;8:443.

Falk S, Gorše I, University of Oxford and Wytham Woods Genome Acquisition Lab et al. The genome sequence of the hawkweed Cheilosia, Cheilosia urbana (Meigen, 1822 Wellcome Open Res 2024a;8:311.

Falk S, Poole O, University of Oxford and Wytham Woods Genome Acquisition Lab et al. The genome sequence of a hoverfly, Cheilosia impressa (Loew, 1840). Wellcome Open Res 2024b;9:74.

Falk S, Poole O, University of Oxford and Wytham Woods Genome Acquisition Lab et al. The genome sequence of the parsley Cheilosia, Cheilosia pagana (Meigen, 1822). Wellcome Open Res 2024c;9:54.

Falk S, University of Oxford and Wytham Woods Genome Acquisition Lab, Natural History Museum Genome Acquisition Lab et al. The genome sequence of the large burdock Cheilosia, Cheilosia vulpina (Meigen, 1822). Wellcome Open Res 2021;6:351.

Falk S, Woodcock KJ, University of Oxford and Wytham Woods Genome Acquisition Lab et al. The genome sequence of the yarrow Cheilosia, Cheilosia vernalis (Fallén, 1817). Wellcome Open Res 2024d;9:188.

Falk S, Woodcock KJ, University of Oxford and Wytham Woods Genome Acquisition Lab et al. The genome sequence of a hoverfly, Cheilosia scutellata (Fallén, 1817). Wellcome Open Res 2024e;9:125.

Feuk L. Inversion variants in the human genome: role in disease and genome architecture. Genome Medicine 2010;2:11.

Ghiurcuta CG, Moret BME. Evaluating synteny for improved comparative studies. Bioinformatics 2014;30:i9–18.

Goel M, Schneeberger K. plotsr: visualizing structural similarities and rearrangements between multiple genomes. Bioinformatics 2022;38:2922–6.

Goel M, Sun H, Jiao W-B et al. SyRI: finding genomic rearrangements and local sequence differences from whole-genome assemblies. Genome Biology 2019;20:277.

Grabherr MG, Russell P, Meyer M et al. Genome-wide synteny through highly sensitive sequence alignment: Satsuma. Bioinformatics 2010;26:1145–51.

Hackl T, Ankenbrand MJ, Adrichem B van. gggenomes: A Grammar of Graphics for Comparative Genomics. 2023.

He W, Yang J, Jing Y et al. NGenomeSyn: an easy-to-use and flexible tool for publication-ready visualization of syntenic relationships across multiple genomes. Bioinformatics 2023;39:btad121.

Howe K, Bateman A, Durbin R. QuickTree: building huge Neighbour-Joining trees of protein sequences. Bioinformatics 2002;18:1546–7.

Krasheninnikova K, Diekhans M, Armstrong J et al. halSynteny: a fast, easy-to-use conserved synteny block construction method for multiple whole-genome alignments. GigaScience 2020;9:giaa047.

Lewin HA, Richards S, Lieberman Aiden E et al. The earth BioGenome project 2020: Starting the clock. Proceedings of the National Academy of Sciences 2022;119:e2115635118.

Liu D, Hunt M, Tsai IJ. Inferring synteny between genome assemblies: a systematic evaluation. BMC Bioinformatics 2018;19:26.

Logsdon GA, Vollger MR, Hsieh P et al. The structure, function and evolution of a complete human chromosome 8. Nature 2021;593:101–7.

Mitchell R, Falk S, Woodcock KJ et al. The genome sequence of a hoverfly, Cheilosia grossa (Fallén, 1817). Wellcome Open Res 2024;9:616.

Quigley S, Damas J, Larkin DM et al. syntenyPlotteR: a user-friendly R package to visualize genome synteny, ideal for both experienced and novice bioinformaticians. Bioinformatics Advances 2023;3:vbad161.

Ried T, Schröck E, Ning Y et al. Chromosome painting: a useful art. Human Molecular Genetics 1998;7:1619–26.

Wang T, Antonacci-Fulton L, Howe K et al. The Human Pangenome Project: a global resource to map genomic diversity. Nature 2022;604:437–46.

Yu G, Smith DK, Zhu H et al. ggtree: an R package for visualization and annotation of phylogenetic trees with their covariates and other associated data. Methods in Ecology and Evolution 2017;8:28–36.

