## Supplementary Material for "ntSynt-viz: Visualizing synteny patterns across multiple genomes"

### Supplementary Information for "ntSynt-viz: Visualizing synteny patterns across multiple genomes"

**Supplementary Table S1.** Descriptions of ntSynt-viz parameters.

| Parameter | Required? | Description | Description of value for parameter (if applicable) | Default value (if applicable) |
|---|---|---|---|---|
| **--blocks** | Y | Synteny blocks TSV | Format output by ntSynt. TSV file with columns: block_id, genome assembly, chromosome, start, end, strand, number of supporting minimizers, reason for discontinuity with previous synteny block. | N/A |
| **--fais** | Y | FAI files for input genomes | Can be a list on the command line, or a file listing the FAI files (one per line). | N/A |
| **--name_conversion** | N | File to define name conversions for final plot | TSV file with columns: old name, new name. The new names cannot have spaces; if you want spaces in your output ribbon plot, all underscores will be converted to spaces at that step. | N/A |
| **--tree** | N | Tree to plot next to the ribbons in the final plot and determine the genome order | Newick file format, with the names matching the new names in the --name_conversion file, if specified. | N/A |
| **--target-genome** | N | Genome to place at the top of the ribbon plot | Genome name | N/A |
| **--normalize** | N | Normalize the strand of the chromosomes relative to the target (top) genome in the ribbon plots | N/A | N/A |
| **--indel** | N | Indel size threshold | Integer (bp) | 50,000 |
| **--length** | N | Minimum synteny block length | Integer (bp) | 100,000 |
| **--seq_length** | N | Minimum chromosome sequence length | Integer (bp) | 500,000 |

| | | | | |
|---|---|---|---|---|
| **--keep** | N | List of chromosomes to show in the visualization | List in the format genome_name:chromosome. All chromosomes with links to these sequences will also be shown. | N/A |
| **--centromeres** | N | File listing coordinates of centromeres, or any other genomic feature | TSV file with headers: bin_id,seq_id,start,end. These correspond to: genome_name (same as new name in --name_conversion if specified), chromosome, start, end. | N/A |
| **--prefix** | N | Prefix for output files | String | ntSynt-viz_ribbon-plot |
| **--format** | N | Output format of ribbon plots | "png" or "pdf" | png |
| **--scale** | N | Length of scale bar | Length in bases | 100e6 |
| **--height** | N | Height of plot | Height in cm | 20 |
| **--width** | N | Width of plot | Width in cm | 50 |
| **--no-arrow** | N | Do not draw arrows | Only used with --normalize; results in no arrows being drawn on the ribbon plot to indicate reverse complementation. | N/A |
| **--ribbon_adjust** | N | Ratio for adjusting spacing beside ribbon plot | Float value. Increase if any labels are cut off, and decrease white space to the left of the ribbon plot. | 0.1 |
| **--force** | N | Force a re-run of the entire pipeline | N/A | N/A |
| **-n** | N | Dry-run for pipeline | N/A | N/A |

**Supplementary Table S2**. Accessions for the human genome assemblies compared using ntSynt and visualized with ntSynt-viz. All accessions are available on NCBI, except for those indicated with an asterisk, which are found on CNCB.

| Human individual | Individual Info | Haploid or diploid? | Accession(s) |
|---|---|---|---|
| **CHM13 (T2T)** | Functionally haploid cell line | Haploid | GCA_009914755.3 |
| **HG002** | Ashkenazi Jew male | Diploid | *Maternal:* GCA_021951015.1<br>*Paternal:* GCA_021950905.1 |
| **CN1** | Han Chinese male | Diploid | *Maternal:* GWHCBHM00000000*<br>*Paternal:* GWHCBHQ00000000* |
| **HuRef** | European male | Diploid | *Prime:* GCA_000212995.1<br>*Alternate:* GCA_000002125.1 |
| **KSA001** | Saudi Arabian female | Diploid | *Maternal:* GCA_037177555.1<br>*Paternal:* GCA_037177635.1 |
| **Han1/HG00621** | Han Chinese male | Haploid | GCA_024586135.1 |
| **HG01243** | Puerto Rican male | Haploid | GCA_018873775.2 |
| **mHomSap3** | Mixed ancestry (African, European, Native American) male | Diploid | *Maternal*: GCA_016695395.2<br>*Paternal:* GCA_016700455.2 |
| **PGP1** | North Eastern European male | Haploid | GCA_020497115.1 |
| **KOREF** | Korean male | Haploid | GCA_020497085.1 |

**Supplementary Table S3**. NCBI accessions for the *Cheilosia* genus genome assemblies compared using ntSynt and visualized with ntSynt-viz and NGenomeSyn.

| Species | NCBI Accession | Number of chromosomes | Genome size (Mbp) |
|---|---|---|---|
| ***Cheilosia grossa*** | GCA_963082955.1 | 6 | 361.5 |
| ***Cheilosia impressa*** | GCA_948293265.1 | 6 | 382.0 |
| ***Cheilosia pagana*** | GCA_936431705.1 | 6 | 353.9 |
| ***Cheilosia scutellata*** | GCA_955612985.1 | 6 | 470.3 |
| ***Cheilosia soror*** | GCA_949372485.1 | 6 | 479.9 |
| ***Cheilosia urbana*** | GCA_946477585.1 | 5 | 545.1 |
| ***Cheilosia variabilis*** | GCA_951230905.1 | 7 | 414.7 |
| ***Cheilosia vernalis*** | GCA_949126925.1 | 6 | 431.8 |
| ***Cheilosia vulpina*** | GCA_916610125.1 | 6 | 404.6 |

**Supplementary Table S4.** ntSynt-viz commands used to generate ribbon plots for the human and hoverfly synteny blocks.

| Dataset | Zoom? | Command |
|---|---|---|
| **Human** | No | plot_gggenomes.py --blocks ntSynt.k24.w1000.synteny_blocks.tsv --fai fais.tsv --name_conversion name_conversions.tsv --target-genome T2T --normalize --centromeres T2T_centromeric_regions.format-for-gggenomes.renamed.bed --haplotypes haplotypes.tsv --indel 10000 --ribbon_adjust 0.15 |
| **Human** | Yes – T2T chromosome 8 | plot_gggenomes.py --blocks ntSynt.k24.w1000.synteny_blocks.tsv --fai fais.tsv --name_conversion name_conversions.tsv --target-genome T2T --normalize --centromeres T2T_centromeric_regions.format-for-gggenomes.renamed.bed --haplotypes haplotypes.tsv --indel 10000 --keep T2T:8 --scale 100e6 --ribbon_adjust 0.15 |
| **Hoverfly** | No | plot_gggenomes.py --blocks ntSynt.k24.w1000.synteny_blocks.tsv --fais fais.tsv --name_conversion cheilosia_asm-species_conversion.renamed.tsv --tree mt_assemblies.aln.fa.renamed.renamed.treefile --normalize --indel 100000 --prefix ntsynt-viz.cheilosia --scale 100e6 --target-genome C._impressa |
| **Hoverfly** | Yes – *C. soror* chromosome OX443561.1 | plot_gggenomes.py --blocks ntSynt.k24.w1000.synteny_blocks.tsv --fais fais.tsv --name_conversion cheilosia_asm-species_conversion.renamed.tsv --tree mt_assemblies.aln.fa.renamed.renamed.treefile --normalize --indel 100000 --prefix ntsynt-viz.cheilosia --scale 100e6 --target-genome C._impressa --keep C._soror:OX443561.1 |

**Supplementary Table S5.** Runtime and peak memory usage for ntSynt-viz and NGenomeSyn runs visualizing all chromosomes.

| Dataset | Tool | Time (min) | Peak memory (GB) |
|---|---|---|---|
| **Human** | ntSynt-viz | 1.90 | 0.41 |
| **Human** | NGenomeSyn | 0.21 | 0.59 |
| ***Cheilosia*** | ntSynt-viz | 1.02 | 0.39 |
| ***Cheilosia*** | NGenomeSyn | 0.11 | 0.32 |

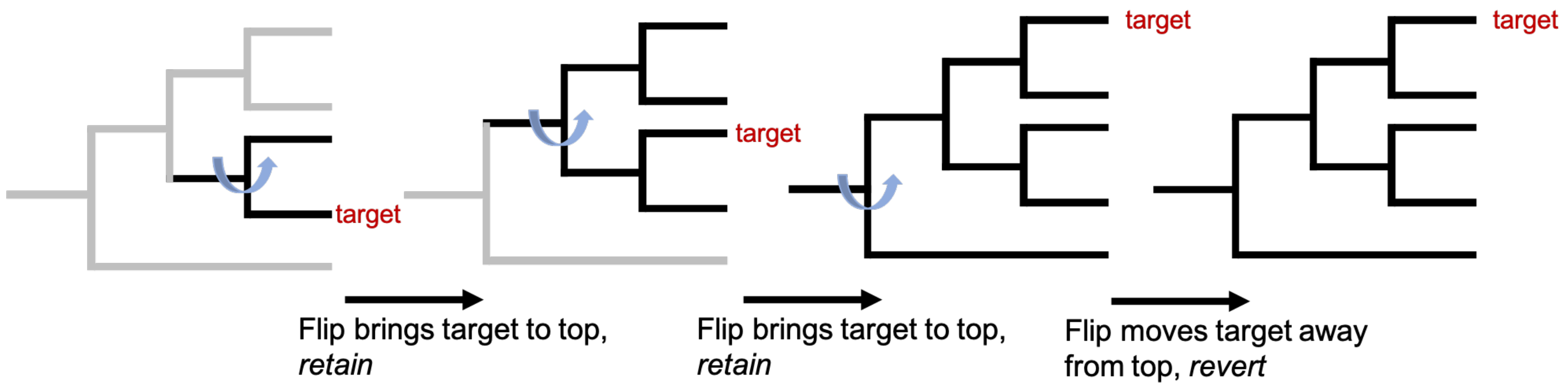

**Supplementary Fig. S1.** ntSynt-viz algorithm for ensuring that the target genome (when --target-genome is specified) is at the top of the cladogram and thus the final ribbon plot. We use rotations around internal nodes of the constructed cladogram to flip the target genome to the top of the cladogram, while retaining the overall tree topology.

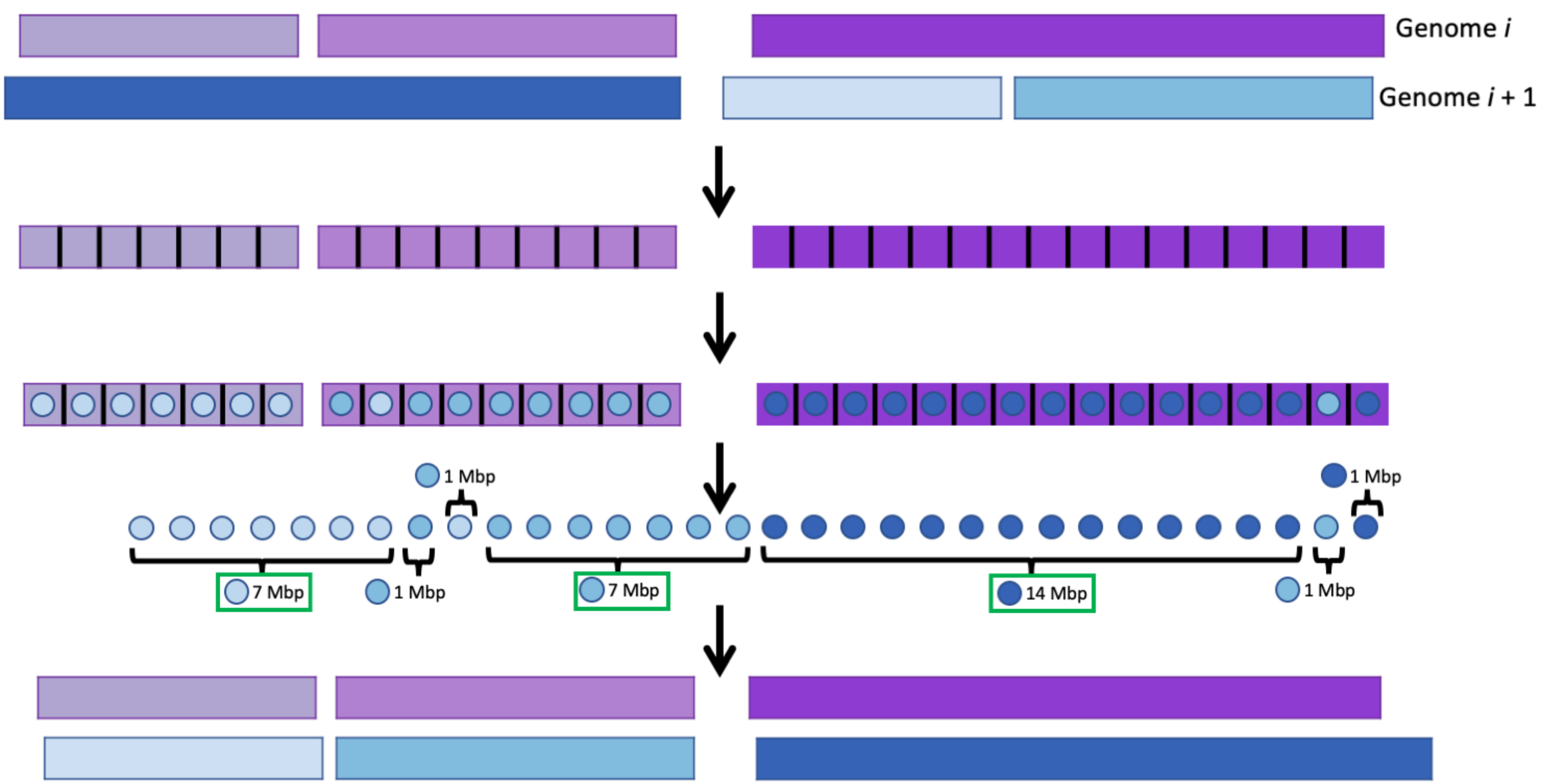

**Supplementary Fig. S2.** ntSynt-viz algorithm for sorting chromosomes. For each pair of ordered genomes, $i$ and $i+1$, the synteny block coordinates between these genomes are compared to determine the ordering of the genome $i+1$ chromosomes. First, the genome $i$ chromosomes are split into non-overlapping tiles (default size 1 Mbp). For each tile, the block lengths for the genome $i+1$ chromosomes are tallied for all synteny blocks that overlap those tile's coordinates. The genome $i+1$ chromosome with the highest length after this tally is assigned to that tile (represented with blue coloured circles). After all tiles are assigned, the assignments are represented as a list, and the run lengths for each chromosome determined. The list is filtered to only contain the longest run length for each $i+1$ chromosome (represented with the green rectangles), and this list represents the ordering of chromosomes for genome $i+1$.

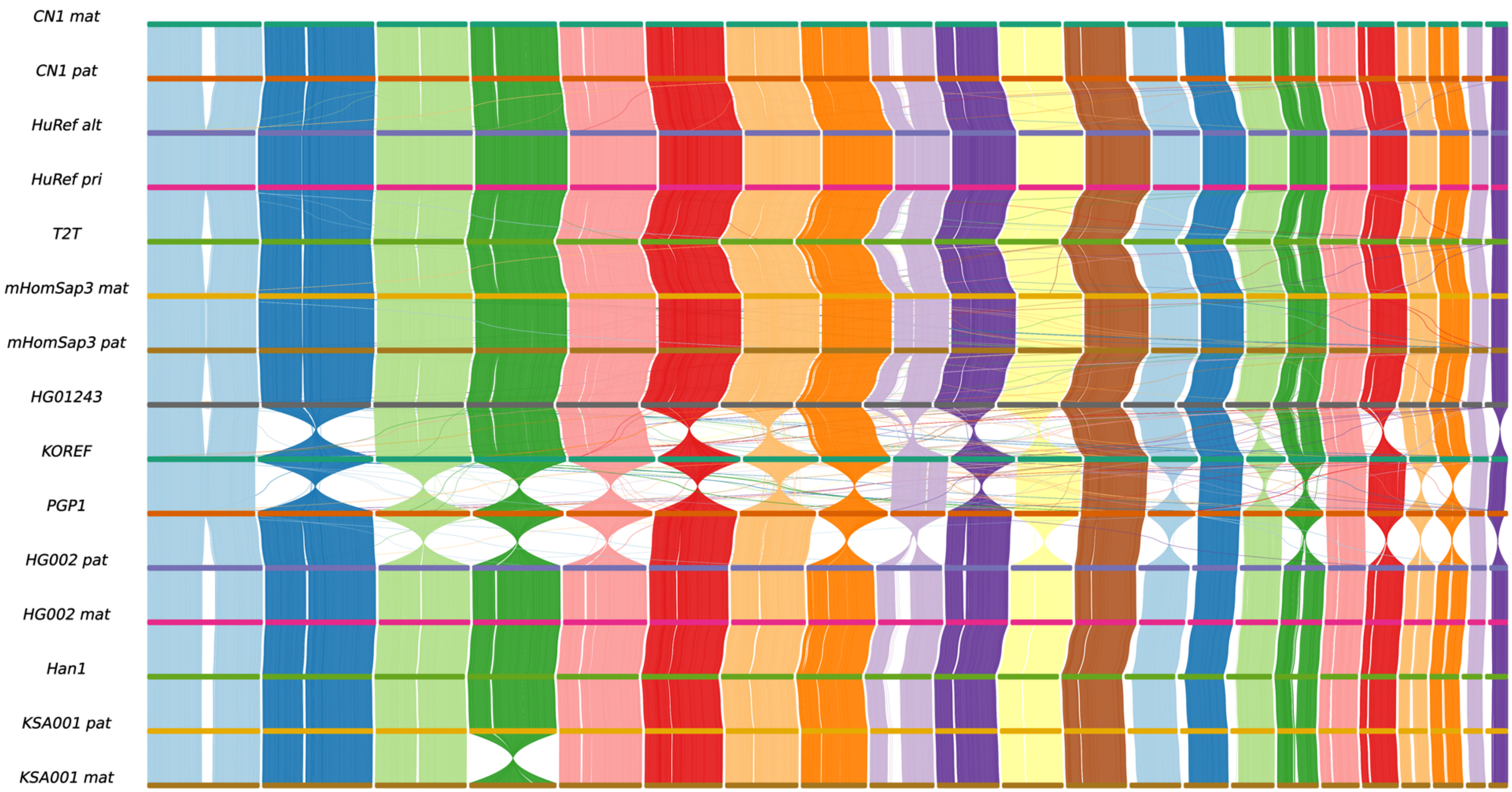

**Supplementary Fig. S3.** NGenomeSyn plot generated using ntSynt synteny blocks between 14 genome sequence assemblies from 10 human individuals.

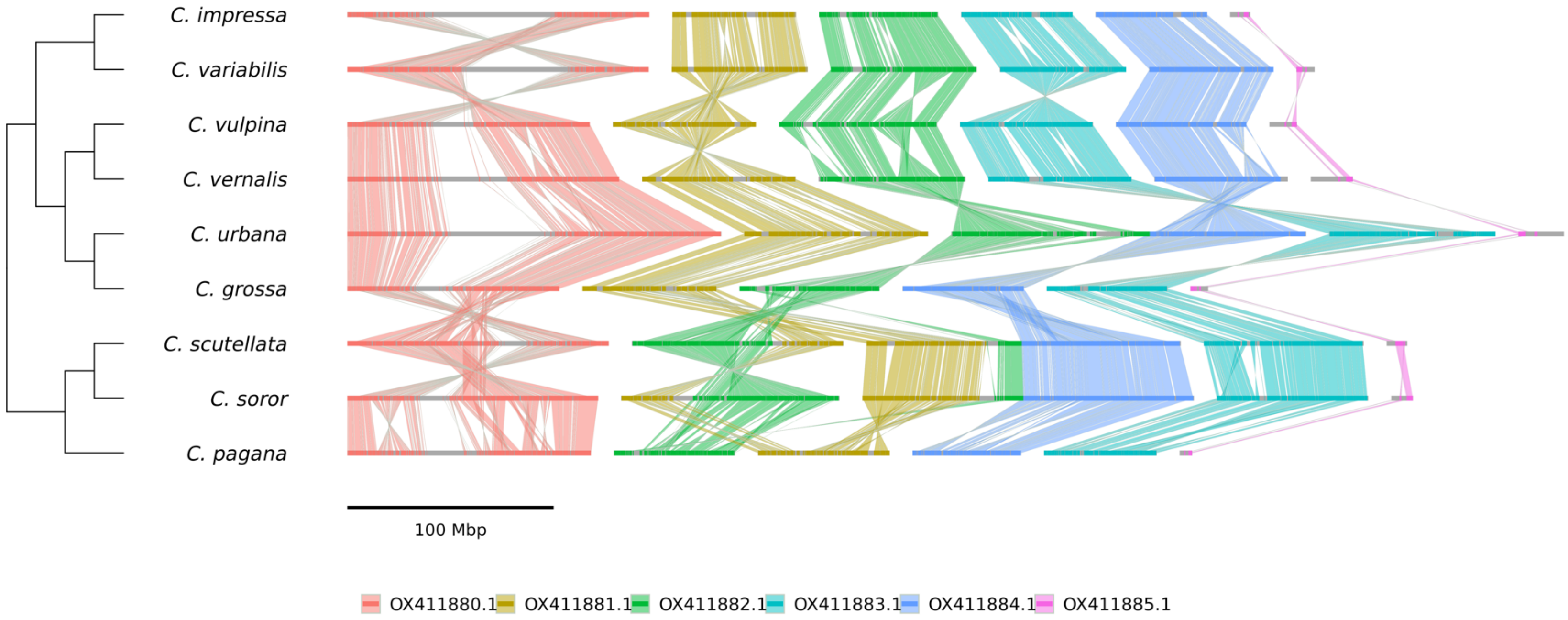

**Supplementary Fig. S4.** ntSynt-viz plot visualizing ntSynt synteny blocks between 9 *Cheilosia* genomes without strand normalization.
